# 3C-seq-captured chromosome conformation of the hyperthermophilic archaeon *Thermofilum adornatum*

**DOI:** 10.1101/2021.04.30.439615

**Authors:** Andrey Sobolev, Mikhail Solovyev, Valeria Ivanova, Tatiana Kochetkova, Alexandr Merkel, Sergey Razin, Sergey Ulianov, Alexander Tyakht

## Abstract

Three-dimensional structure of chromosomes displays diverse patterns across the tree of life, with compartments, interaction domains and loops being quite universally observed. The archaeal kingdom remains understudied to this extent so far, despite representing an interesting area from evolutionary and other perspectives.

Here we describe the spatial chromosomal organization of a hyperthermophilic crenarchaeon *Thermofilum adornatum* strain 1910b based on high-throughput chromosome conformation capture (3C-seq) approach. The chromosome contact map showed a curved secondary diagonal almost orthogonal to the main one. No evidence of chromosome loops was present. We were able to identify boundaries of different strengths between chromosome interaction domains (CIDs) albeit moderate. The plaid-like patterns previously reported for *Sulfolobus* archaea were not observed. However, the calculation of A/B compartments divided the genome into 2 domains that were different by the density of predicted highly expressed genes and location of origins.

Further comparison of these domains with whole-genome gene expression profiles will allow to test whether these domains represent expression-associated compartments. If so, it is possible that they represent primitive compartments evolutionarily older than the plaid patterns of *Sulfolobus* and higher eukaryotes. Further exploration of 3D chromatin in all branches of archaeal diversity will elucidate the evolution of the links between structural and functional organization in live organisms.

## Introduction

Archaeal chromosomes are often found in bacteria-like condensed nucleoid structures, with various proteins playing a role in its organization. However, their nucleoid-associated proteins (NAPs) and mechanisms of DNA compactization closely resemble those of Eukaryotes (Bell and White 2010). From all three kingdoms of life, Archaea stand out for sure since they are able to efficiently combine different mechanisms of chromatin organization from Bacteria and Eukaryotes for adaptation to severe environmental conditions such as high salt concentrations, acidic pH and high temperatures (Laursen, Bowerman, and Luger 2021).

Similarly to Eukaryotes and Bacteria, archaeal chromosomes are organized at multiple levels of compactization, with many proteins responsible for these levels, namely histone-like and nucleoid-associated proteins such as Alba family proteins. Moreover, proteins of structural maintenance of chromosomes (SMC) family are also involved in shaping archaeal DNA. Almost all *Crenarchaeota* species also include Cren7 protein, which monomers bind DNA in a head-to-tail fashion while structuring the DNA into an S-shaped filament (Laursen, Bowerman, and Luger 2021). *Thermofilum pendens* Hrk5 is a notable exception since this crenarchaeal species does not encode Cren7 homologs (Guo et al. 2008). Sul7d, another SH3-domain protein which is known to be specific for the *Sulfolobus* family, coexists with Cren7, constituting about 5% of total protein in the cell, therefore, it is classified as abundant chromatin architectural protein involved in DNA kinking (Laursen, Bowerman, and Luger 2021).

Histone-like proteins of Archaea share a common ancestor with eukaryotic core histones (Mattiroli et al. 2017). However, their amino acid structure differs from the Eukaryotes since most archaeal histones lack N-terminal end crucial for histone modifications. Segregation of chromatin segments with different sets of histone modifications facilitate formation of chromosomal compartments (Bian et al. 2020). Noteworthy, several species from the Asgard superphylum along with *Thermofilum pendens Hrk5* were reported to include an N-terminal tail in their histone protein sequences (Henneman et al. 2018; Laursen, Bowerman, and Luger 2021), suggesting that due to their possible DNA-binding properties they can participate in compartmentalization and/or formation of hypernucleosomes. In addition, reported compartmentalization in *Sulfolobus* species lacking histones implies that a different mechanism of spatially segregating chromosome segments is possible.

However, organization of the chromosome structure at the histone level varies within archaeal phyla. For instance, *Methanothermus fervidus*, a euryarchaeal species, contains histone-like proteins HMfA and HMfB. The latter tends to dimerize in solution but forms tetrameric tertiary structure in presence of DNA. *Thermococcus kodakarensis* histones form nucleosome-like structures of different size (usually three to five dimers of histone-like proteins), around which DNA is wrapped in a spiral-like manner (Laursen, Bowerman, and Luger 2021). In general, size of these structures can vary depending on the ability to form a “hypernucleosome” that is likely determined by dimer–dimer interactions as well as stacking interactions between individual layers of the hypernucleosome (Henneman et al. 2018). In Crenarchaeota, histones and histone-like proteins are generally absent. Nevertheless, *Thermofilum pendens* Hrk5, along with species from *Vulcanisaeta* and *Caldivirga*, were reported to encode such proteins (Henneman et al. 2018). However, *Vulcanisaeta* and *Caldivirga* encode Cren7 in addition to the histone proteins (Peeters et al. 2015). This limited dispersion of histone homologs and Cren7 in archaeal genomes suggests that histones and Cren7 may serve redundant roles (Laursen, Bowerman, and Luger 2021). histones are known to be involved in chromosome compartmentalization.

Along with the presence of hypernucleosomes, archaeal genomes are also reported to be organized into loops and structures resembling chromosomal interaction domains (CIDs) in Bacteria. Using high-throughput chromosome conformation capture approach (Hi-C), such CID-like domains were observed not only in *Euryarchaeota* (Cockram C, Thierry A, Gorlas A, Lestini R, Koszul R 2020), but also in crenarchaeal genomes (Takemata and Bell 2021) of *Sulfolobus* spp., where they are suggested to be formed as a result of locally strong transcriptional processes. In addition, they are facilitated by a *Sulfolobus*-specific protein termed coalescin (ClsN) which was earlier reported to be involved in the formation of A/B compartments in these Archaea (Takemata, Samson, and Bell 2019). These compartments are associated with transcription in *Sulfolobus*, however, in case of *Euryarchaeota* such bipartition of the chromosome is of structural nature only and is not likely associated with functional differences (Cockram C, Thierry A, Gorlas A, Lestini R, Koszul R 2020). These studies emphasize the diversity of chromosome organization in archaeal genomes since these microorganisms share bacterial and eukaryotic properties.

Hyperthermophilic microorganisms are characterized by the ability to grow optimally at temperatures ≥80°C. Most of them belong to the Archaea domain. They are extremely interesting objects for study not only from the point of view of the evolution of life (Gribaldo and Brochier-Armanet 2006; Nasir, Kim, and Caetano-Anollés 2014), ecology (Huber, Huber, and Stetter 2000) and biotechnology (Han et al. 2019), but also from the point of view of adaptation of the primary functions of the cell (replication, transcription, translation) to life at such high temperatures. Description of spatial chromosome organization in hyperthermophilic archaea will further elucidate the mechanisms of their tolerance to harsh environmental conditions. We applied the high-throughput chromosome conformation capture technique 3C-seq to evaluate its structure in the *Thermofilum adornatum* strain 1910b (optimal growth temperature: 92°C) belonging to the *Thermoproteales* order (Zayulina et al. 2020).

## Methods

### Cultivation

The strain 1910b^T^ was cultivated (0.6 l) at optimal growth conditions on strictly anaerobic modified Pfennig medium (Podosokorskaya et al. 2011) under N_2_ in the gas phase at 80°C and pH 5.75, supplemented with 0.1 g/l of yeast extract, 1/100 (v/v) of sterile culture broth filtrate of *Desulfurococcus* sp. 1910a and 1.0 g/l of glucose as the substrate. The grown cells were collected at an early stationary phase by centrifugation at 17,000 x *g* for 20 min.

### 3C-seq library preparation and sequencing

The cell culture (5-7×10^7^ cells/ml) was cooled to room temperature for 10 min. The cells were pelleted from the growth media, washed twice with growth media without yeast extract, and fixed in growth media without yeast extract with 3% formaldehyde for 15 min with occasional mixing. The reaction was stopped by the addition of 2 M glycine to give a final concentration of 125 mM. Cells were centrifuged (17,000 × *g*, 10 min, 4 °C), resuspended in 50 μl of 1× PBS, snap-frozen in liquid nitrogen and stored at −80 °C. Defrozen cells were disrupted using FastPrep-24 5G homogenizer and Lysing Matrix A (MP Biomedicals, USA) and additionally lysed in 1 ml isotonic buffer (50 mM Tris-HCl pH 8.0, 150 mM NaCl, 0.5% (v/v) NP-40 substitute (Fluka), 1% (v/v) Triton-X100 (Sigma), 1× Halt™ Protease Inhibitor Cocktail (Thermo Scientific) on ice for 15 min. Cells were centrifuged at 20,000 × g for 5 min at 4 °C, resuspended in 200 μl of 1× NEBuffer 2 (NEB), and pelleted again. The pellet was resuspended in 200 μl of 0.3% SDS in 1× NEBuffer 2 and incubated at 37 °C for 1 hour. Then, cells were centrifuged at 20,000 x g for 5 min at 4°C, washed with 200 μl of 1× NEBuffer 2 and resuspended in 1× CutSmart buffer (NEB) supplemented with 1% of Triton X-100 (Sigma). One hundred U of HpaII enzyme (NEB) were added, and the DNA was digested overnight (14–16 hours) at 37 °C with shaking (1,400 rpm). On the following day, additional 100 U of HpaII enzyme were added, and the cells were incubated for an additional 2 hours. HpaII was then inactivated by incubation at 65 °C for 20 min. After HpaII inactivation, the cells were harvested for 10 min at 20,000 x g, washed with 300 μl of 1× T4 DNA ligase buffer (Fermentas), and resuspended in 300 μl of 1× T4 DNA ligase buffer. Cohesive DNA ends were ligated in the presence of 75 U of T4 DNA ligase (Fermentas) at 16 °C for 4 hours. The cross-links were reversed by overnight incubation at 65 °C in the presence of proteinase K (1 μg/μl) (Sigma) and 0.5% of SDS. After cross-link reversal, the DNA was purified by single phenol-chloroform extraction followed by ethanol precipitation with 20 μg/ml glycogen (Thermo Scientific) as the co-precipitator. After precipitation, the pellets were dissolved in 100 μl 10 mM Tris-HCl pH 8.0. To remove residual RNA, samples were treated with 50 μg of RNase A (Thermo Scientific) for 45 min at 37 °C. To remove residual salts and DTT, the DNA was additionally purified using Agencourt AMPure XP beads (Beckman Coulter). The DNA was then dissolved in 500 μl of sonication buffer (50 mM Tris-HCl (pH 8.0), 10 mM EDTA, 0.1% SDS) and sheared to a size of approximately 100–1,000 bp using a VirSonic 100 (VerTis). The samples were concentrated (and simultaneously purified) using AMICON Ultra Centrifugal Filter Units to a total volume of approximately 50 μl. The DNA ends were repaired by adding 62.5 μl MQ water, 14 μl of 10× T4 DNA ligase reaction buffer (Fermentas), 3.5 μl of 10 mM dNTP mix (Fermentas), 5 μl of 3 U/μl T4 DNA polymerase (NEB), 5 μl of 10 U/μl T4 polynucleotide kinase (NEB), 1 μl of 5 U/μl Klenow DNA polymerase (NEB), and then incubating at 20 °C for 30 min. The DNA was purified with Agencourt AMPure XP beads and eluted with 127 μl of 10 mM Tris-HCl (pH 8.0). To perform an A-tailing reaction, the DNA samples were supplemented with 15 μl 10× NEBuffer 2, 3 μl of 10 mM dATP, and 4.5 μl of 5 U/μl Klenow (exo-) (NEB). The reactions were carried out for 30 min at 37 °C in a PCR machine, and the enzyme was then heat-inactivated by incubation at 65 °C for 20 min. The DNA was purified using Agencourt AMPure XP beads and eluted with 100 μl of 10 mM Tris-HCl (pH 8.0). Illumina TruSeq adapters were ligated by adding 12 μl 10× T4 DNA ligase reaction buffer (Fermentas), 6 μl of Illumina TruSeq adapters and 2 μl of 5 U/μl T4 DNA ligase (Fermentas). Adapter ligation was performed at 22 °C overnight. DNA was then purified with Agencourt AMPure XP beads and eluted with 30 μl of 10 mM Tris-HCl (pH 8.0). Test PCR reactions containing 5 μl of the samples were performed to determine the optimal number of PCR cycles required to generate sufficient PCR products for sequencing. The PCR reactions were performed using KAPA High Fidelity DNA Polymerase (KAPA) and Illumina PE1.0 and PE2.0 PCR primers (10 pmol each). The temperature profile was 5 min at 98 °C, followed by 6, 9, 12, 15, and 18 cycles of 20 s at 98 °C, 15 s at 65 °C, and 20 s at 72 °C. The PCR reactions were separated on a 2% agarose gel containing ethidium bromide, and the number of PCR cycles necessary to obtain a sufficient amount of DNA was determined based on the visual inspection of gels (typically 10-12 cycles). Four preparative PCR reactions were performed for each sample. The PCR mixtures were combined, and the DNA was purified using Agencourt AMPure XP beads and eluted with 50 μl of 10 mM Tris-HCl (pH 8.0).

The sequencing was performed on Illumina HiSeq platform in 2 × 150 bp reads format.

### Data analysis

The sequences were preprocessed using BBMap tool suite (https://sourceforge.net/projects/bbmap/). The reference genome of *T. adornatum* strain 1910b (GenBank: CP006646.1) was downloaded from the NCBI Genbank (Dominova et al. 2013). The chromosome contact maps were obtained using Juicer v1.6 (Durand et al. 2016) and verified with hiclib (Imakaev et al. 2012). The raw contact map demonstrated two secondary diagonals. In fact, due to the circular nature of the archaeal genome, these diagonals would fuse into one when the genome is shifted (see Supplementary Fig. 1). Therefore, we moved the first 200000 nucleotides of the genome sequence provided by GenBank to the end of the sequence. The borders between chromosomal interaction domains (CIDs) were identified using the insulation score method (Crane et al. 2015) implemented in *cooltools* v0.3.2; the optimal window size was determined empirically as 80 Kbp. Additionally, we used the *detect* function of *chromosight* v1.4.1 to detect CIDs. Analysis of compartmentalization was performed using *cooltools* v0.3.2 as described previously (Lieberman-Aiden et al. 2009). The predicted highly expressed (PHX) genes were identified with EMBOSS v6.6.0 (Rice, Longden, and Bleasby 2000) utilities *cusp* and *cai* using the previously published PHX of *T. pendens* (Anderson et al. 2008) as the training set. We considered genes with codon adaptation index in the top 5% as PHX. Gene convergence profile was constructed as described before (Mizuguchi et al. 2014). In brief, each 1 Kbp bin was assigned with an orientation score depending on the orientation of genes in it: 1 in case of prevalence of downstream genes, −1 for the prevalence of upstream genes and 0 otherwise. For the convergence profile, the 1 Kbp convergence score was calculated as the weighted sum of positive gene orientation bins 50Kbp upstream and negative bins 50Kbp downstream. Then, a 5 Kbp convergence profile was constructed by averaging 1 Kbp convergence score in each five non-overlapping bins. For the 5 Kbp orientation profile, orientation scores in each five non-overlapping bins were averaged. The whole-chromosome 3D form was reconstructed and visualized using GenomeFlow v2.0 (Trieu et al. 2019) (LorDG algorithm), with additional use of EVR for reconstruction (Hua and Ma 2019) and MeshLab for 3D visualization.

Identification of putative replication origins was performed using the Z-curve approach implemented in the Ori-Finder2 (Luo, Zhang, and Gao 2014) tool searching for *Sulfolobaceae* motifs with the p-value cutoff of 10^-6^. All other parameters were left as defaults. The search for the genes encoding putative DNA replication proteins, SMC-like and histone-like proteins in *T. adornatum* genome was carried out using blastp at the NCBI website. We used the coverage cutoff of 80% and E-value cutoff of 0.0001 were considered homological. For the SMC-like proteins, the sequence of *T. pendens* SMC domain proteins (GenBank ID: ABL77933.1 and ABL78969.1) were used as a query for blastp. For the histone-like proteins, we used *T. pendens Hrk5* histone (ABL77757.1) as a query. For the Alba proteins, the sequences of *T. pendens* Alba proteins (ABL77621.1, ABL77941.1 and ABL78659.1) were used. Finally, we performed a search for the homologs of ClsN protein (ADX84150.1) discovered previously (Takemata, Samson, and Bell 2019). We investigated each sequence of interest using hmmscan against Pfam database.

## Results

As the result of 3C-seq, 62.5 mln read pairs alignable to the *T. adornatum* genome were produced, among them - 12.8 mln read pairs representing valid 3-C/Hi-C contacts. The resulting chromosome contact map confirmed the validity of the experimental part; the coverage was sufficient to provide the resolution as high as 3 Kbp. Interestingly, the map manifested patterns not commonly observed in previous archaeal studies using 3C-seq/Hi-C approaches: the main diagonal was complemented with an almost orthogonal curved secondary diagonal (following a cyclic shift of the reference genome and the contact map, respectively; see Methods) (Fig. 1A). Similar curved diagonals reflecting asymmetric juxtaposition of chromosome arms have been observed in contact maps of *Bacillus subtilis* with parS sites inserted at ectopic chromosomal positions (Wang et al. 2017, 2015). The originality of the chromosome conformation is also suggested by its 3D reconstruction (Fig. 1B). Interestingly, in addition to the rod-shaped structure that was expected to be found according to the contact map, we observed a rotation of this structure in a spiral-like manner. Moreover, a visible difference was noticed in the juxtaposed areas of the chromosome — the left half (as on Fig. 1B) seemed to be more compacted - reflecting the fact that the chromosome arms are closer to each other than in the lower half, according to the contact map (Fig. 1A).

**Fig. 1.**
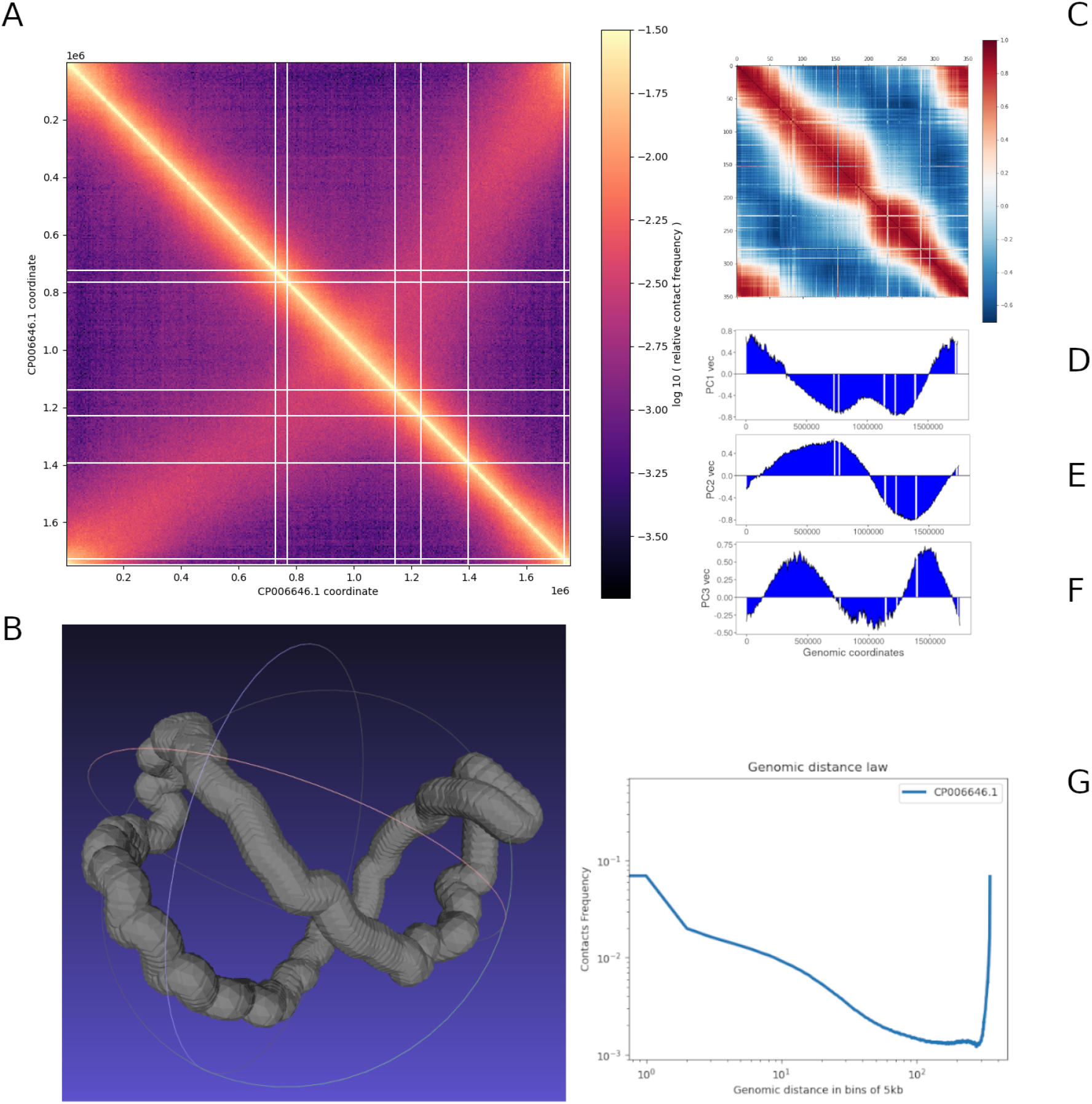
Spatial organization of *Thermofilum adornatum* 1910b chromosome. **(A)** Chromosome contact map (5 Kbp resolution). **(B)** Reconstruction of 3D structure of chromosome with EVR. **(C)** Pearson’s correlation matrix with the first 3 principal components (**(D), (E)** and **(F)**, respectively) derived from A-B compartment analysis. **(G)** Scaling curve.

### Compartmentalization analysis indicates possible bipartition of the genome into two different-sized domains

Interestingly, we did not observe “plaid”-like patterns on the contact map previously reported for the two hyperthermophilic archaea species: *Sulfolobus acidocaldarius* and *S. islandicus* from the same *Crenarchaeota* phylum (Takemata, Samson, and Bell 2019). Formal application of the methodology for identifying the compartments resulted in various separations of the genome into domains of different size (Fig. 1D, 1E, 1F). Due to the orthogonality of two first principal components, we assumed that they represent chromosome division into domains along two orthogonal axes, with the second principal component possibly reflecting the functional differences in the chromosome areas. The second principal component changes its sign at ~1.02 Mbp (in the circularly shifted genome, see Methods), and this area corresponds to the intersection point of the main and secondary diagonals observed on the contact map; moreover, there is a visible thinning in the correlation map at this region (Fig. 1C). We considered the domain with positive eigenvector values as a domain I of total length ~900 Kbp (a small region of ~55 Kbp length with positive eigenvector value was also included in this domain). The second domain - with negative eigenvector values, along with the small structure on the left - was considered as domain II of total length ~795 Kbp (Fig. 1B).

To check if the two identified genomic domains could be considered as the most primitive case of transcriptionally active A/B compartments (“2 x 2 checkerboard”), it is necessary to assess the gene expression along the *T. adornatum* chromosome. Due to the lack of the respective experimental data, we resorted to an *in silico* prediction of the highly expressed genes (PHX) based on codon usage bias (Karlin et al. 2005). Using the data for a closely related species *T. pendens* (Anderson et al. 2008) as a training set, we identified 95 predicted highly expressed genes and compared their location with the results of compartmentalization analysis. Interestingly, we found that the number of PHX was significantly different between domains (binomial test, p = 0.007313): 61 (~64.2%) of the PHX corresponded to domain I, and 34 (~35.8%) of the PHX belonged to domain II (Fig. 2A). We also compared GC content in the domains - since the regions with higher expression levels are known to be characterized by higher GC content (Du et al. 2018) - but found no significant differences (Mann-Whitney test, p > 0.05) (Fig. 2B). No significant difference was also observed between GC content in the whole genome and in bins that contain PHX (Mann-Whitney test, p > 0.05) (Fig. 2C).

**Fig. 2.**
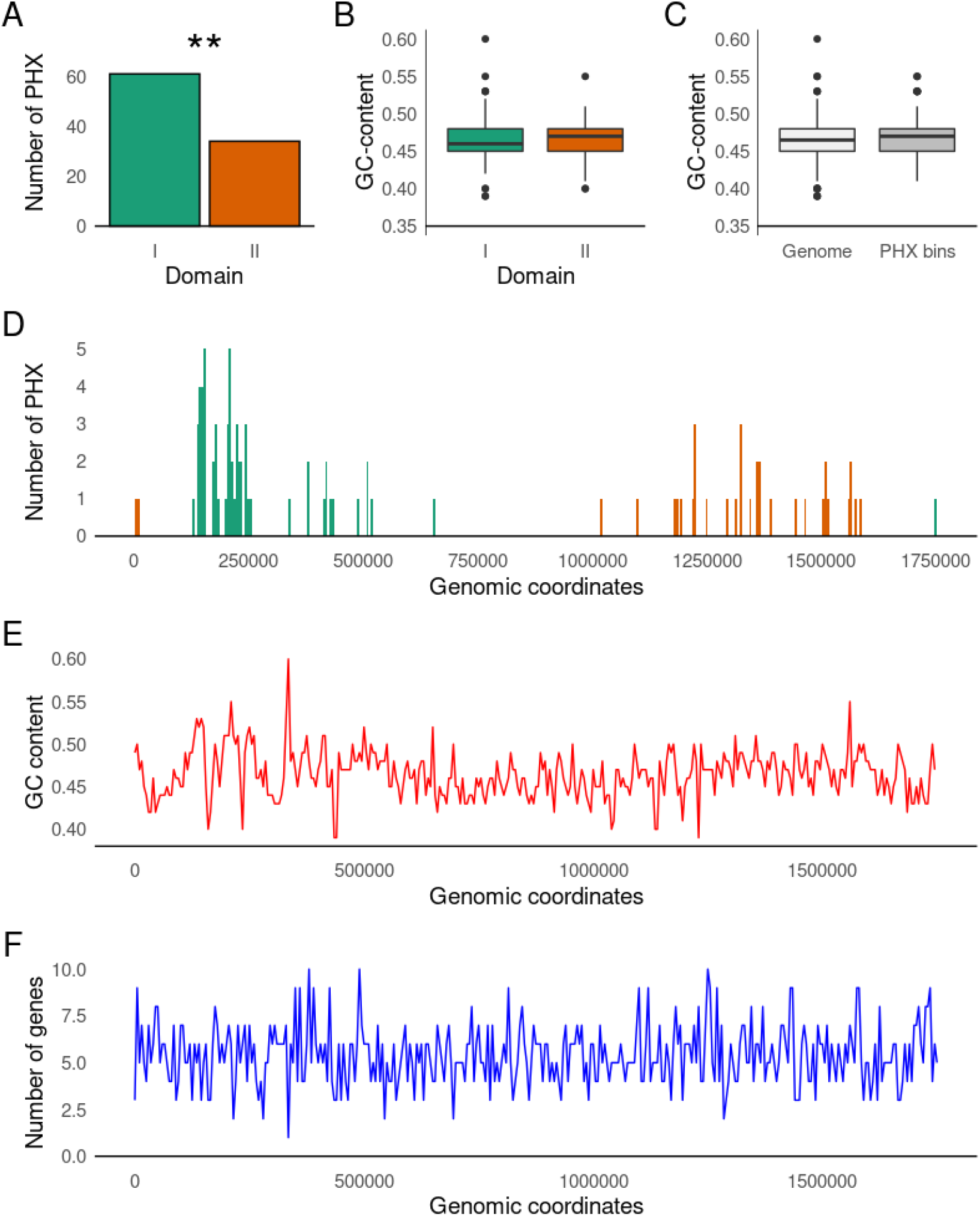
Genomic features compared with the 3C-seq data. **(A)** Number of PHX genes in the domains (p = 0.007313, binomial test). **(B)** GC content in the domains (p > 0.05, Mann-Whitney U-test). **(C)** Whole-genome GC-content compared to GC content in bins containing PHX (p > 0.05, Mann-Whitney U-test). **(D)** Distribution of predicted highly expressed genes along the genome. **(E)** GC content and **(F)** gene density of the chromosome calculated per 5 Kbp bins.

Analysis of the distribution of PHX genes along each domain showed that 46 (48.4%) of them formed a large genomic cluster in the region between ~0.13 and ~0.25 Mbp (Fig. 2D). Mostly, this cluster consists of unannotated proteins, ribosomal proteins, genes involved in transcription and various polymerases subunits. We further checked if the presence of this cluster was associated with an increased gene density (Fig. 2E) or with higher GC-content (Fig. 2F) because of the possible gene density bias and GC-content bias. Neither significant increase of gene density in the region of the characterized cluster (p = 0.1293, Mann-Whitney U-test) nor correlation between the GC-content and number of genes per bin (p = 0.5542, Spearman’s r = 0.0318) were observed. Interestingly, regions corresponding to the intersection of main and secondary diagonal (750-1000 Kb and 1700-100 Kb, Supplementary Fig. 1) were depleted of PHX which indicates the decrease of transcriptional activity in these regions.

In *Sulfolobus* species, all three origins of replication are located in the compartment displaying higher overall transcriptional activity. We investigated the distribution of predicted origin(s) of replication along the *T. adornatum* genome.

### Links of replication origins with the 3D genome of *Thermofilum*

Unlike the bacterial genomes, those of Archaea often contain multiple origins of replication (Lundgren et al. 2004; Pelve et al. 2012; Robinson and Bell 2007). The proximity of genes to the origin of replication is associated with higher expression levels in bacteria (Couturier and Rocha 2006) and in archaea (as shown in *Sulfolobus* having multiple origins (Flynn et al. 2010)). The origin sequences in Archaea are located near the initiator genes, such as *Cdc6/Orc1* (Wu et al. 2014). These proteins can carry out functions of both *Cdc6* and *Orc1* homologous proteins in Eukaryotes. Similarly to the Eukaryotes, replication origins of Archaea are usually flanked with origin recognition boxes (ORB). Nevertheless, origins of replication display wide diversity in the set of initiator downstream genes and ORB elements (Pelve et al. 2013; Wu et al. 2012; Majerník and Chong 2008). Moreover, some Archaea are able to replicate their chromosome even without using replication origins (Kelman and Kelman 2018).

There are several approaches available to predict origin(s) location in the genome sequence but it is necessary to take into account that GC skew in archaeal chromosomes can be absent in the case of multiple replication origins (Arakawa, Suzuki, and Tomita 2009). For this reason, we applied the Z-curve approach implemented in Ori-Finder2 (Luo, Zhang, and Gao 2014), a tool designed specifically for the archaeal genomes. We searched for the *Sulfolobaceae* motifs to identify putative origin(s) of replication (Fig. 3D). As a result, four origins of replication were detected (Table 1, Fig. 3B, Supplementary Material 1). For two of them, located at 0.33 Mbp and 0.52 Mbp, homologous origins in the DOriC database (Luo and Gao 2019) were found - AORI10010294 from *Haloarcula* CBA1115 (*Euryarchaeota*) and AORI10010034 - from *Thermofilum pendens* Hrk5. As the latter species is closely related to *T. adornatum*, we suggest there is higher evidence that the second prediction is true.

**Fig. 3.**
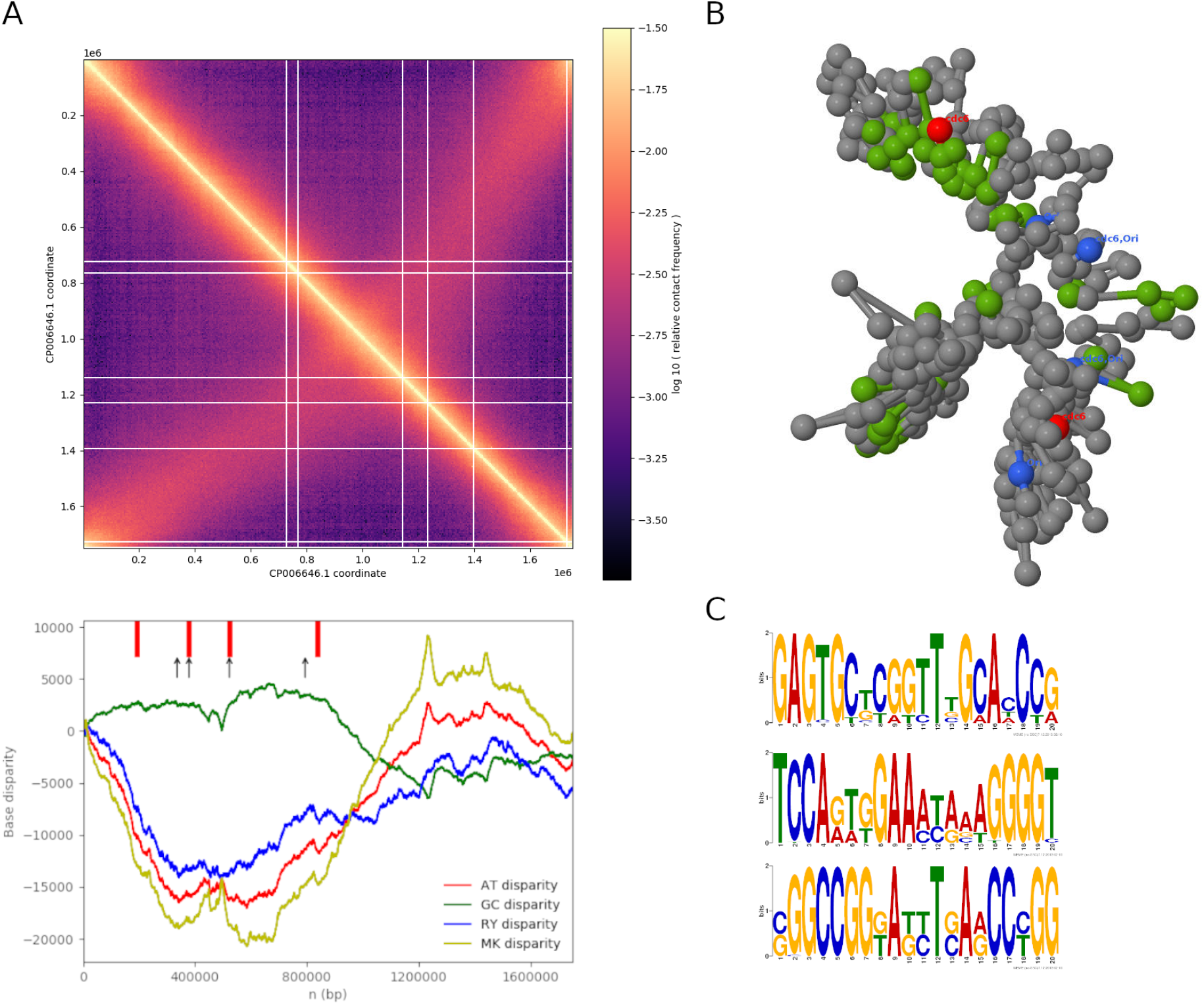
Distribution of replication genes, origins and PHX in the *T. adornatum* chromosome. **(A)** Contact map (5 Kbp resolution) compared to Ori-Finder2 results. Red lines indicate the position of detected genes of the replication proteins. Black arrows indicate the position of the replication origins detected by Ori-Finder2. **(C)** Whole-genome 3D structure reconstructed using the LorDG algorithm (end view). Red, green and blue regions indicate the position of genes of replication, origins of replication and PHX, respectively (some colocalized). **(D)** *Sulfolobaceae* motifs used for detection of replication origin in Ori-Finder2.

**Table 1.** Origins of replication detected using Ori-Finder2. (†) — genomic coordinates in the shifted genome. NA — not available.

| Start | End | Strand | GC content | DOriC ID | Number of detected ORBs |
| --- | --- | --- | --- | --- | --- |
| 536553<br>(336553†) | 538457<br>(338457†) | + | 62.55% | AORI10010294 | 4 |
| 578794<br>(378794†) | 579192<br>(379192†) | + | 48.99% | NA | 3 |
| 723312<br>(523312†) | 723614<br>(523614†) | + | 44.37% | AORI10010034 | 2 |
| 994517<br>(794517†) | 994785<br>(794785†) | + | 45.52% | NA | 3 |

Four DNA replication genes were detected in the genome at the following regions: 0.19, 0.37, 0.52 and 0.83 Mbp in the shifted genome (Fig. 3C) (the respective coordinates in the original GenBank annotation are 0.39, 0.57, 0.72 and 1.03 Mbp, see Table 2). We investigated these genes using hmmscan against Pfam database and blastp against *Crenarchaeota* genomes. As a result, each of the sequences AGT34835.1 and AGT35192.1 had CDC6 C-terminal winged helix and AAA ATPase domain that were distinctive for Cdc6/Orc1 homologs in archaea. The sequence AGT35513.1 included an AAA domain only. Finally, AGT35033.1 had two helix-turn-helix domains related to bacterial regulatory proteins of the arsR family, therefore we considered it a false positive finding unrelated to the replication process. As for the blastp search, all high-similarity matches displayed results similar to the Pfam search, with all the sequences except for AGT35033.1 corresponding to Cdc6/Orc1 homologs and/or AAA family ATPases. Replication origins detected by Ori-Finder2 and genes of replication proteins were often adjacent, but some genes were not annotated as Cdc6/Orc1 homologs in our case (Table 2). These findings indicate that *T. adornatum* 1910b genome contains at least 3 Cdc6/Orc1 homologs with putative replication origins located nearby.

**Table 2.** DNA replication genes in *Thermofilum adornatum* 1910b genome and their functions. (†) — genomic coordinates in a shifted genome. (*) and (f) — region note and protein function according to the GenBank annotation, respectively.

| Location in GenBank annotation | Strand | Accession ID | Comments |
| --- | --- | --- | --- |
| 394645 - 395950<br>(194645 - 195950†) | + | AGT34835.1 | Hypothetical protein<sup>f</sup>. ORC1-type DNA replication protein; PRK00411* |
| 579192 - 579516<br>(379192 - 379516†) | - | AGT35033.1 | Hypothetical protein<sup>f</sup>. Arsenical Resistance Operon Repressor and similar prokaryotic, metal regulated homodimeric repressors. ARSR subfamily of helix-turn-helix bacterial transcription regulatory proteins* |
| 723614 - 724847<br>(523614 - 524847†) | + | AGT35192.1 | Cell division control protein 6; Reviewed; PRK00411<sup>f</sup> |
| 1039802 - 1040930<br>(839802 - 840930†) | - | AGT35513.1 | Hypothetical protein<sup>f</sup>. Cdc6-related protein, AAA superfamily ATPase (Replication, recombination and repair); COG1474* |

Another way to confirm these findings is to assess gene expression around the origins of replication. Here, the cluster of PHX at the 0.13 — 0.25 Mbp region that has been identified earlier is noteworthy: one of the detected replication genes falls nearby this region. Despite the absence of replication origins in this area, the replication gene located at 0.19 Mbp belongs to Cdc6/Orc1 homologs according to the annotation. Presence of cluster of PHX, in addition to the two local minima in GC content in this area (Fig. 2) and Cdc6/Orc1 homolog located at 0.19 Mbp, might indicate that this region can likely contain an origin of replication as well. However, PHX is not the ultimate way to assess gene expression, and thorough investigation of the transcriptional activity in domains requires an RNA sequencing analysis.

Finally, origins of replication, genes of the replication proteins and PHX profile were mapped to the constructed whole-genome 3D structure (Fig. 3C). Clearly, all predicted replication origins, along with genes of replication proteins and cluster of PHX described above, were found to be colocalized in one half of the genome if it is hypothetically divided in half in a longitudinal manner. (Interestingly, none of the origins were at the cross of the main and secondary diagonal - unlike other known prokaryotic cases when the secondary diagonal is present.) These results likely confirm the assumption that compartmentalization analysis results represent bisection of *T. adornatum* 1910b chromosome in two orthogonal directions, possibly making one chromosome arm more active than other.

### Detection of chromosomal loops and interaction domains (CIDs)

According to the 3D structure of chromosome, *T. adornatum* genome does not include any loop structures. The obtained contact map also did not show any features that could be resembling loops. Formal analysis did not show any of these structures neither via HICCUPS nor via Chromosight (despite the fact that both algorithms performed well on the published crenarchaeal data (Takemata and Bell 2021)). Recent studies of *Sulfolobus* species indicate that the loop formation in these Archaea is correlated with the location of ribosomal genes — locations of loop anchors seem to be enriched with such genes (Takemata and Bell 2021). However, our findings suggest that *Thermofilum* species organize their genomes in other ways.

At first sight, the map did not show pronounced square-like structures along the diagonal resembling chromosomal interaction domains (CIDs). However, closer examination of the main diagonal region and whole-genome reconstruction of chromosome form (Fig. 1C, Fig. 3C) suggests that such structures may be present in the *T*. *adornatum* chromosome. For a more formal analysis, we applied insulation score (Crane et al. 2015) - one of the CID identification algorithms that performed best for prokaryotes in a benchmark study (Magnitov et al. 2020) - using the window size of 80 kbp - to yield 35 CID boundaries (Fig. 4E). Some of the detected borders coincided with the ones observed visually upon closer examination of the main diagonal. Boundary re-analysis using Chromosight produced results similar to the case of insulation score (Fig. 4D), resulting in 31 boundaries.

**Fig. 4.**
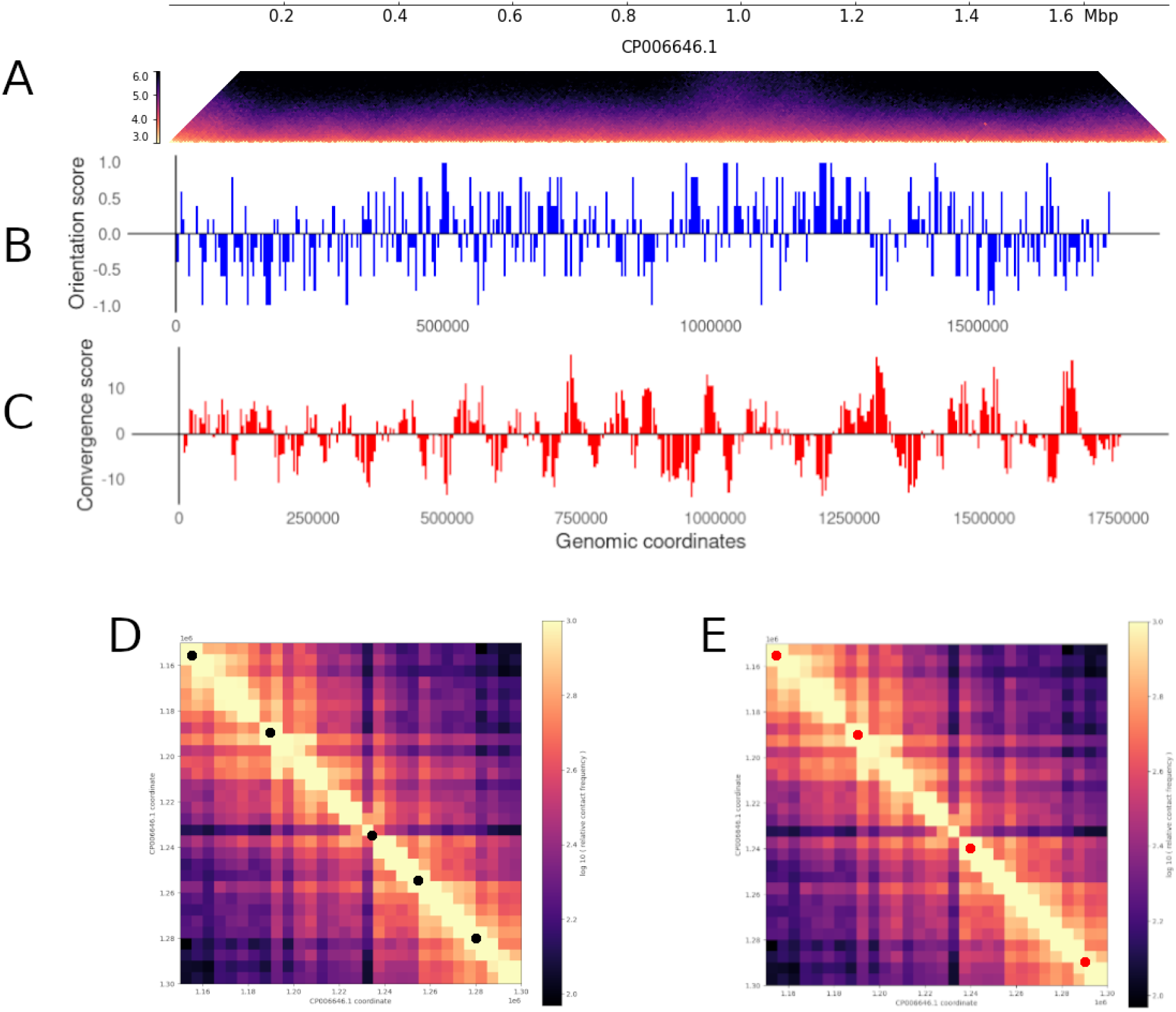
Gene convergence bias and CID boundaries detection results. **(A)** Diagonal-adjacent part of the contact map. **(B)** Gene orientation and **(C)** convergence profiles calculated at 5 Kbp resolution. **(D)** Fragment of the contact matrix (from 1.15 Mbp to 1.3 Mb) with CID boundaries detected with Chromosight. **(E)** The same fragment with CID boundaries detected with insulation score (window size = 80 Kb). Black dots on **(D)** and red dots on **(E)** represent inter-CID boundaries detected by Chromosight and insulation score.

### Search for SMC

Proteins of the SMC superfamily are known to be widely involved in shaping the chromosome structure. This superfamily is highly conserved across different kingdoms of the tree of life including Archaea, with various proteins carrying out a wide variety of functions including DNA recombination, reparation, juxtaposition of chromosome arms and formation of chromatin loops (Hassler, Shaltiel, and Haering 2018). However, some of the eukaryotic SMC superfamily members are absent in some Archaea. Instead, SMC complexes of weak structural homology carry out the same functions. For instance, recent study reports that the cohesin is absent in *Sulfolobus* species of *Crenarchaeota* (Takemata and Bell 2020) and the CIDs boundaries are formed mainly due to high level of transcription along with the activity of SMC-like protein coalescin (ClsN), which is also believed to be responsible for the formation of A/B-compartments, displaying an enrichment in transcriptionally inactive genome compartment of these species (Takemata, Samson, and Bell 2019). We looked for SMC homologs of *T. pendens* Hrk5 and ClsN homologs of *Sulfolobus* spp. in the *T. adornatum* 1910b genome using blastp. In the case of *T. pendens*, both query sequences corresponded to the SMC family ATPases in *T. adornatum* sequence (E-value = 0.0), although with low coverage (47.03% and 42.28%, respectively). The blastp search for the ClsN homologs did not yield significant matches. Apparently, the SMC superfamily members are present in *T. adornatum* but the microorganism might be using structurally different proteins to maintain the chromosome structure.

Along with the SMC proteins activity, gene transcription can play a role in chromosome ordering. In the genome of *Saccharomyces pombe* yeast, cohesin loading sites are correlated with convergent gene regions (Mizuguchi et al. 2014). We constructed profiles of gene orientation and convergence as described in Mizuguchi et al. (Figs. 4B, C; see Methods) to check if the “direction bias” and/or gene convergence bias is present in the genome and if it is related to the location of the detected origins of replication, since cohesin-loading sites are located nearby OriC (Guillou et al. 2010). We then compared the resulting profile with insulation score profile, distribution of detected inter-CID boundaries and location of replication proteins and origins (Supplementary Fig. 2). Interestingly, we observed a good visual concordance between these features. Replication proteins and/or origins corresponded to the inter-CID boundaries and changes of convergence score sign. These observations indicate that *T. adornatum* might utilize the insulation principles similar to those of the Eukaryotes.

### Presence of histone-like proteins

Similarly to the Eukaryotes, Archaea are known to possess histone proteins involved in compaction and functional organization of their genomes. In some species, however, the histones are not present. Such organisms contain histone-like proteins such as homologues of bacterial DNA benders HU or nucleoid-associated proteins from the Alba family (Peeters et al. 2015), forming multimer structures of different size termed hypernucleosomes that wrap the DNA around it in a left-hand manner. However, known archaeal histones are absent in most *Crenarchaeota* (Henneman et al. 2018). To investigate the presence of histone proteins in *T. adornatum* 1910b, we used the known *T. pendens* Hrk5 histone as a query for BLAST search. We found that one of the hypothetical *T. adornatum* 1910b proteins (ABL77757.1) and the aforementioned histone shared 75.58% of amino acid sequence (E-value = 8e-46). In addition, one of the blastp hits corresponded to the 52 amino acids long hypothetical protein of the different *T. adornatum* strain 1505 (identity = 76.92%, E-value = 1e-24). Moreover, Alba proteins are also present in this genome annotated as DNA-binding (AGT35146.1, 88.17% aa identity) or hypothetical (AGT36222.1, 77.78% aa identity) proteins. Presence of this putative histone and Alba proteins in *T. adornatum* 1910b indicates that this archaeon can contain hypernucleosomes, which distinguishes it, along with *T. pendens* Hrk5, *Caldivirga maquilingensis* and *Vulcanisaeta distributa*, from most investigated *Crenarchaeota* species (Henneman et al. 2018).

## Discussion and conclusions

Archaeal chromosomes, like the bacterial and eukaryotic ones, display a wide variety of patterns at different levels of genome organization, namely hypernucleosomes, compartments, chromatin loops and chromosomal interaction domains. Here, we described the structural and functional organization of *Thermofilum adornatum* 1910b chromosome. Its peculiar features include a structure resembling a twisted loop, with some regions more adjacent to each other and organized in more sophisticated spindle structures, taking the form of a helix. Recent studies described three-dimensional chromosomal structure for a pilot set of *Crenarchaea* and *Euryarchaea* (Takemata, Samson, and Bell 2019; Cockram C, Thierry A, Gorlas A, Lestini R, Koszul R 2020) of *Sulfolobus, Haloferax, Thermococcus* and *Halobacterium* species. The *Thermofilum* genus belongs to the *Thermoproteales* order in *Crenarchaeota* which is the earliest branch in the phylum and remains understudied. Hence, genome organization in this order could be expected to differ considerably from one of *Sulfolobales*. Our findings confirm this statement, since *Thermofilum* contact map lacks plaid pattern and chromatin loops described in *Sulfolobus*, even at the relatively high resolution of 5 Kbp that we were able to obtain. Therefore, our results show that the 3D genome organization can vary drastically also within the same archaeal phylum. This being said, there are several signs suggesting possible chromatin compartmentalization in the *Thermofilum*. For instance, all detected replication origins and DNA replication genes found with Ori-Finder2, along with the predicted cluster of PHX around 0.13 — 0.25 Mbp, are colocalized in the domain which we suggested to be transcriptionally active (domain I). Moreover, the PHX distribution turned out to be different between the domains, with the enrichment in domain I. It is tempting to speculate the two domains represent primitive compartments evolutionarily older than the plaid patterns manifested by *Sulfolobus* and higher eukaryotes.

In bacteria, highly expressed genes are known to be clustered near the replication origin (Couturier and Rocha 2006). In *T. adornatum*, we did not observe a strong correlation between PHX and location of replication origins, despite the colocalization of PHX cluster with one of the Cdc6/Orc1 homologs. Therefore, experimental validation of the PHX is required to estimate the possible correlation between the gene expression and localization of replication origins in *T. adornatum*. Moreover, the detected origins require experimental validation, too, since none of the algorithms designed for the search of replication origins have achieved 100% precision yet (Sernova and Gelfand 2008). For instance, the replication genes detected by Ori-Finder2 and considered one of the identified Cdc6/Orc1 homologs as false positive because this gene was not known to be involved in the DNA replication process in Archaea. However, two of the detected origins had been annotated in the DOriC database, with one of them corresponding to the origin of *T. pendens* Hrk5. Lastly, the fact that all detected origins and Cdc6/Orc1 homologs are colocalized in the domain I supports our hypothesis about variable transcriptional activity between the 2 characterized domains. To confirm this hypothesis, it is necessary to compare our findings with the RNA-seq data. For further investigation, analysis of gene expression at different growth phases and under different stress conditions (e.g., amended growth media and/or temperature, presence of actinomycin D) will be insight-gaining. Examining transcriptional changes in *T. adornatum* and how they are related to the chromatin organization will help us to validate our results and clarify whether gene expression is related to the compartment-like domains in the genome of interest, as the sole presence of such domains has been recently shown to not necessarily be related to the transcriptional activity (Cockram C, Thierry A, Gorlas A, Lestini R, Koszul R 2020).

Recent studies showed that *Sulfolobus* species utilize a specific protein termed coalescin (ClsN) to maintain the chromosomal B compartment (as opposed to a more active A compartment) and facilitate CIDs bundling in it (Takemata and Bell 2021). In our case, the *T. adornatum* lacks ClsN homologs indicating that this archaeon might utilize other proteins of the SMC superfamily. This is supported by presence of at least two SMC proteins with distant structural resemblance of their homologs in *T. pendens*. At least, Alba proteins (AGT35146.1, AGT36222.1) likely shape the structure of chromosome at the levels of CIDs and/or compartment-like domains, while the histone-like protein (AGT34931.1) possibly facilitates in formation of hypernucleosomes. The observed colocalization of convergent gene regions, replication origins, inter-CID boundaries and replication genes indicate that *T. adornatum* likely possesses mechanisms of shaping the chromosome structure resembling those described in the Eukaryotes. Complementary methods like ChIP-seq for these proteins might help to elucidate their roles.

A distinctive characteristic of species of *Thermoproteales* order is their cell shape. In particular, members of this order have a form of long rod. This shape is rather peculiar, since other related Crenarchaeota such as members of *Desulfurococcales, Fervidicoccales* or *Sulfolobales* orders manifest regular, irregular or lobed cocci-shaped cells. This elongated and narrow (~0.2 μm) form of the *Thermofilum adornatum* cell can be possibly linked to the chromosome shape we observed - hallmarked with the presence of secondary diagonal on the contact map we have obtained not observed in other Archaea to date. It should be further investigated whether the chromosome is partially or fully stretched along the cell using methods like fluorescent *in situ* hybridization (FISH) to visualize DNA localization inside the cell. Further Hi-C studies of closely related Archaea of this genus such as *T. uzonense* or *T. pendens* will help to check if this peculiar chromosome organization pattern is present in other species of *Thermoproteales*.

Our results show that chromatin architecture at different levels of its organization (and, possibly, its relation to gene expression) varies dramatically between and within orders of Archaea. Further studies of a large representative set covering diverse phylogenetic branches of Archaea are required to elucidate the evolution of 3D genome structure in this kingdom.

