## Supplementary Materials for "3C-seq-captured chromosome conformation of the hyperthermophilic archaeon *Thermofilum adornatum*"

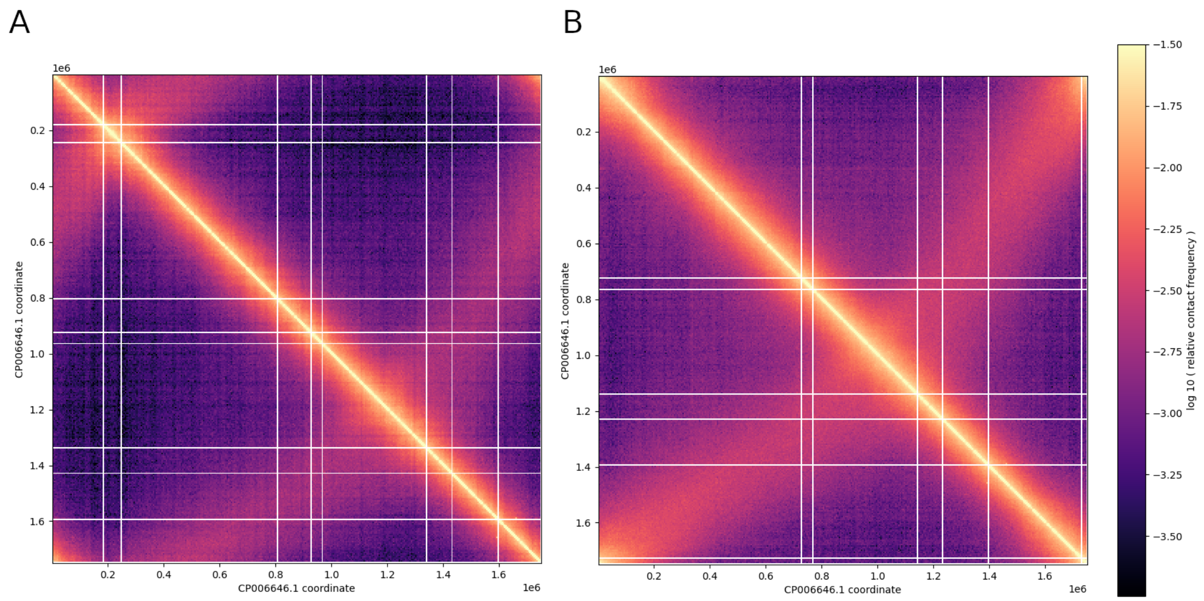


**Supplementary Fig. 1. Circular shift of the contact map.** 5 Kbp contact matrix before **(A)** and after **(B)** circular shift of the genome.

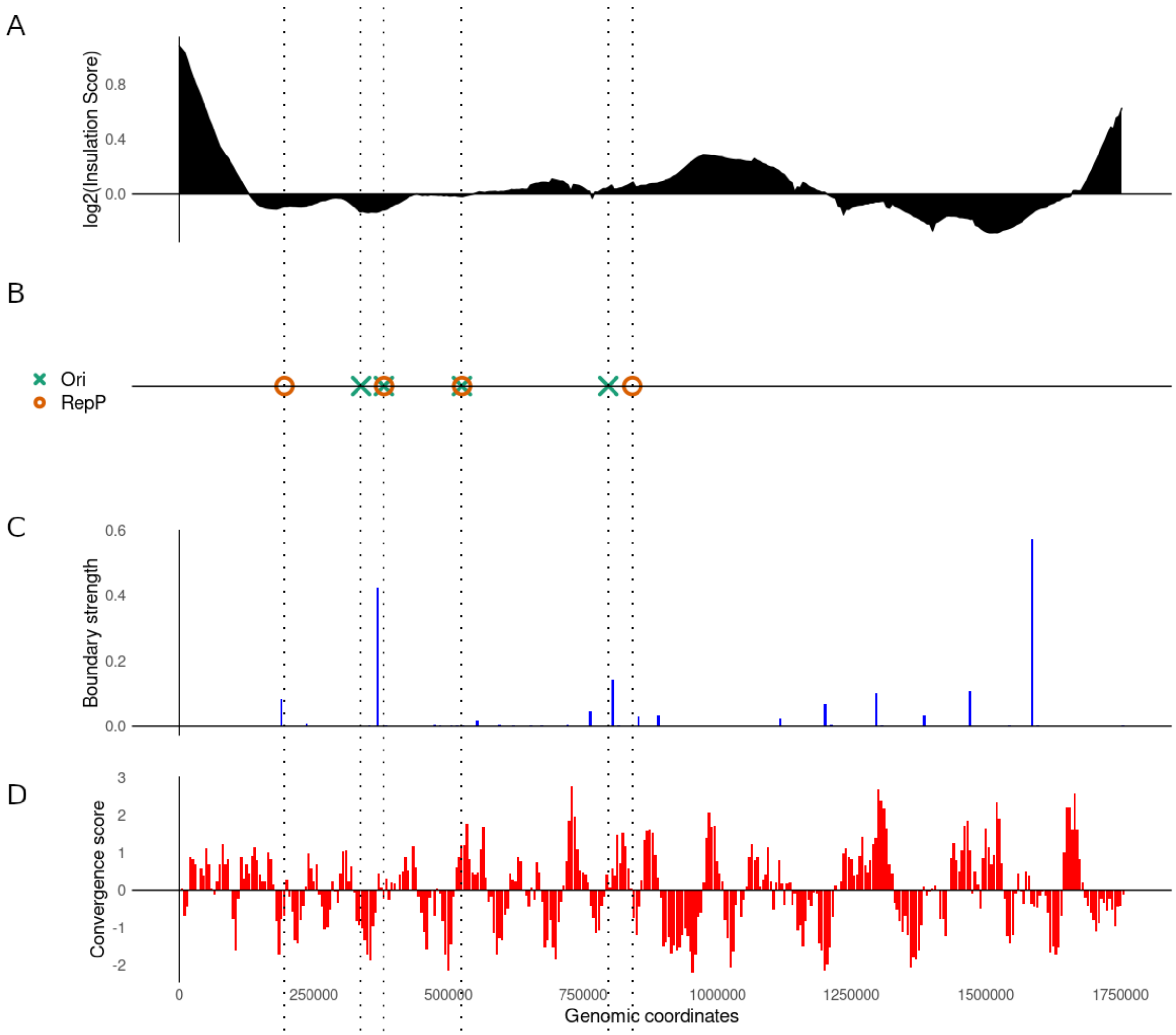


**Supplementary Fig. 2. Insulation tendency in *T. adornatum* genome. (A)** Insulation score profile (window size = 80 Kb). **(B)** Location of replication proteins and origins. **(C)** Inter-CIDs boundary strength. **(D)** Gene convergence profile. Dotted lines on all plots are added for more convenient comparison.

### Supplementary Note 1. Sequences and additional information on replication origins detected by Ori-Finder2.

**Origin 1.** Location: 336554 - 338458 (+). Sequence:
cttaacgctgtctaccggtgtaagcttggtctgcacactctctgtgcggcgctcatattctcccactggtaccccagtgggagttcacgcccttcacccgg
aaacgctctggtttccgggttgcatctcggtaagcatgaaaagtgaggcacggtgtgcgtttctcgaccggatcgaaaggaggtgatccggccgca
ggttcccctacggccaccttgttacgacttctccccccttgggaggcccagattcgacctcccacctcgtcggcaggaggcctcacctgggcctcccT
CGGGTGGAGCGACGGGCGGtgtgtgcaaggagcagggacgtattcaccgcgcgttgatgacacgcggttactagggattccaggttc
acgagggcgagttgcagccctcgattcctactgaggcggggtttaagggattgcctcccccctttcggggtcggaacccgttgtccccgccattgcagc
tcgcgtgtagcccgggggtttcggggcatactgacctgccgtagcccccaccttcctccggcttatcgccggcagtccccctagagtgccccggtccc
gaaggaccgggtagcaactaggggcgggggtctcgctcgttccctgacttaacaggacacctcacggcacgagccggcgacggccatgcacct
cctctcagctcgtctggcaaggtcgttagcctggccttcatcctgctgtcgcccccggtaaggtttccggcgttgacTCCAATTAAACCGCAA
GCTTcacccccttgtggtgctcccccgccaattcctttaagtttcagccttgcggccgtactccccaggcggcaggcttaacggcttccctacggcact

aggcgggcacgaagcccgcctaacacctagcctgcatcgtttacagctgggactacgggggtatctaatcccctttgctcccccagctttcgtccctc
accgtcgggCGCGTGCTAGCCGGGCGCCTtcgccactggtggtcctcctgggattataggatttcgcccctaccccaggagtaccccc
ggcctctcccgccccctagcccggcagtatccctccgctcccacggttgagccgtgggctttaaggagggacttaccgggccggctacggacgcttt
aggcccaataaacgtcccgaccactcgcggggttggtattaccgcggcggctgacaccaagcttgcccccgcttattCGCCCGGCTTTTT
AGACCGGgcaaaagccaccctcatcgggtggcactcggggtgactccgtcacggtttcccgcattgcggaagtttcgcccctggtgcaccccgt
agggcctgggcccttgtctcagtgcccatctcggggctcccgctctcacggcccctacccgttataggcttggcgggccgttaccccgccaactacct
gataggccgcagccccatcctcgggcggctacggcagggatggtgccgtagccctttcggcgtagacccattccagaagtctacacctatggggg
attatccccagtttcccggggttatccccctcccgagggtaggttagccacgtgttactgagccgtccgccggagccggccgtgcccccggctcctag
acttccatgggttagccccaccccgatagcggtcgggtccggcaggatcaaccggagtctggccgcgtgcggaggagtcgataagcttacgcaca
ccgtgcctcactttcttatccccgagagttccgagctagcgctcgtttttctcggggacgcactgtgtagccatgggcggaggccgccttttcattgggag
aggttgcctctcccgttatctaggcgagccgccggtgcccatggcggcaaaattattggttaatggggttgtttataaattttacta

The information on ORB:

| Pattern name | Start | End | Score | P value | Matched sequence |
|---|---|---|---|---|---|
| Sulfolobaceae_motif1 | 294 | 314 | 1 | 1.19E-5 | TCGGGTGGAGCGACGGGCGG |
| Sulfolobaceae_motif1 | 1223 | 1243 | -1 | 1.69E-5 | CCGGTCTAAAAAGCCGGGCG |
| Sulfolobaceae_motif2 | 748 | 768 | -1 | 5.73E-5 | AAGCTTGCGGTTTAATTGGA |
| Sulfolobaceae_motif3 | 969 | 989 | 1 | 8.03E-5 | CGCGTGCTAGCCGGGCGCCT |

**Origin 2.** Location: 378795 - 379193 (+). Sequence:
tagatggcacagggcaacatagcacgcgtcaaaaataggtggaaagaatacttctacctcggcttctatacacgcggcgtactaaaactagcaga
aaagcgcgcgccactttttcaaaacaggctggtaagcatattaggacttgtagacactttgccctcgcacaaaaaatataaacccagaacaaaa
cataatgaagcgagcattagagacttaaacctcgggtcGCCGGGGTAGCTCAGCCTGGgagagcatccggctgaagaacctcgca
agcgtgcgtagacaccggagggtcCCGGGTTCAAGTCCCGGCCCcggcaccattcttgtaaaggtgtttccaaaggttatacgatcta
gtagaggatTGGATTCTAGCCGAGAATGCatgtt

The information on ORB:

| Pattern name | Start | End | Score | P value | Matched sequence |
|---|---|---|---|---|---|
| Sulfolobaceae_motif1 | 229 | 249 | -1 | 1.4E-6 | CCAGGCTGAGCTACCCCGGC |
| Sulfolobaceae_motif1 | 300 | 320 | 1 | 1.31E-10 | CCGGGTTCAAGTCCCGGCCC |
| Sulfolobaceae_motif3 | 373 | 393 | 1 | 7.85E-5 | TGGATTCTAGCCGAGAATGC |

**Origin 3.** Location: 523313 - 523615 (+). Sequence:
aactagaaagggtccttgaaaatcaaagaaaatcctggtgcccCGGCCGGGATTTGAACCCGGgatccttggctcgaaaggccaat
atcctgtcggccgtgcaggctgaCCGGGCTAGACGACCGGGGCttcaagtaggagaggcaaaaaataaaatgaacctaagatttaa
aaacttttcatttggcctatgacaagaacttgtctctacaggtcttctaggggtgacgggaacaacgtataactactgtattttaaatgtacaatgaacgtg
cccaagtttttgtgtaagtttttctata

The information on ORB:

| Pattern name | Start | End | Score | P value | Matched sequence |
|---|---|---|---|---|---|
| Sulfolobaceae_motif1 | 43 | 63 | -1 | 7.52E-12 | CCGGGTTCAAATCCCGGCCG |
| Sulfolobaceae_motif1 | 109 | 129 | 1 | 9.45E-6 | CCGGGCTAGACGACCGGGGC |

**Origin 4.** Location: 794518 - 794786 (+). Sequence:
acatgatacatggaaacgtgtaaaataaaacgtttacgtaaggcaTACAGTGGAACCAGAATGGTggacCGGCCGGGATTC
GAACCCGGgaccaccCGCATGCCAAGCGGACATCCtcccactagacgaccggcccccgttttatgtttagaatactagagatttta
aatagttttcgtgctcttgtgccggttgagtggataaattgttctaatgtgggctccgatgcgcgaaatttatttttgggtactttcatgaatgaaaccat

The information on ORB:

| Pattern name | Start | End | Score | P value | Matched sequence |
|---|---|---|---|---|---|
| Sulfolobaceae_motif1 | 69 | 89 | -1 | 4.63E-10 | CCGGGTTCGAATCCCGGCCG |
| Sulfolobaceae_motif2 | 45 | 65 | -1 | 2.31E-6 | ACCATTCTGGTTCCACTGTA |
| Sulfolobaceae_motif3 | 96 | 116 | 1 | 9.36E-5 | CGCATGCCAAGCGGACATCC |
